# Challenges and Strategies for Recruiting Type 1 Diabetes Families in Kuwait with Strong Beliefs in Familism

**DOI:** 10.1101/570945

**Authors:** Zahra Rahme, Nehad Taha, Hidaia Abdalla, Smitha Abraham, Asma Alhubail, Hala AlSanae, Maria AlMahdi, Fahed Aljaser, Majda Abdelrasoul, Mona Al Khawari, Mohamed Jahromi

## Abstract

Type 1 diabetes (T1D) is one of the most common endocrine and metabolic conditions in children. In fact, this disease in children and adolescents has been increasing exponentially, with Kuwait being ranked second highest in the world regarding the number of T1D incidences. Kuwait is an oil-rich country known for its strong sense of familism, affiliative obedience, and filial obligation. Therefore, a familial study of this disease may disclose certain causative agents responsible for passing the disease on to subsequent generations.

To recruit T1D patients and their family members, three different scenarios were developed. First, since Kuwaiti families are generally obedient to their doctors, the authors decided to recruit the patients through their endocrine physicians. Second, home visits were performed for meeting the families’ requirements. In this case, a team consisting of one nurse, two phlebotomists (a male and a female, since some refused to be seen by the opposite gender), and a driver of the institute’s car was arranged. Finally, two diabetes educators were employed to resolve any issues raised during the recruitment process. Utilizing these approaches helped convince the culturally and religiously oriented Kuwaiti families to participate in this study. In this case, the doctors and educators were not only aware of the obstacles in this population but also sensitive to the families’ beliefs. This paper reports on our experience in recruitment and presents a roadmap for any future familial studies on culturally tailored societies i.e. Arab populations.

## 1. Introduction

Diabetes has reached epidemic proportions throughout the world, thus affecting millions of people. In fact, for every adult diagnosed with diabetes, there is another who is undiagnosed, since chronically elevated blood glucose levels do not often result in symptoms [1]. In Kuwait, there has been a rapid increase in childhood-onset ofT1D incidences over the last several decades, with the current number reported as 41/100,000/year, which is the second highest in the world after Finland [2] (see Figure 1). Given the prevalence of this disease, special actions for preventing and reducing its impact in Kuwait should be one of the highest priorities for scientific research and healthcare.

**Figure 1:**
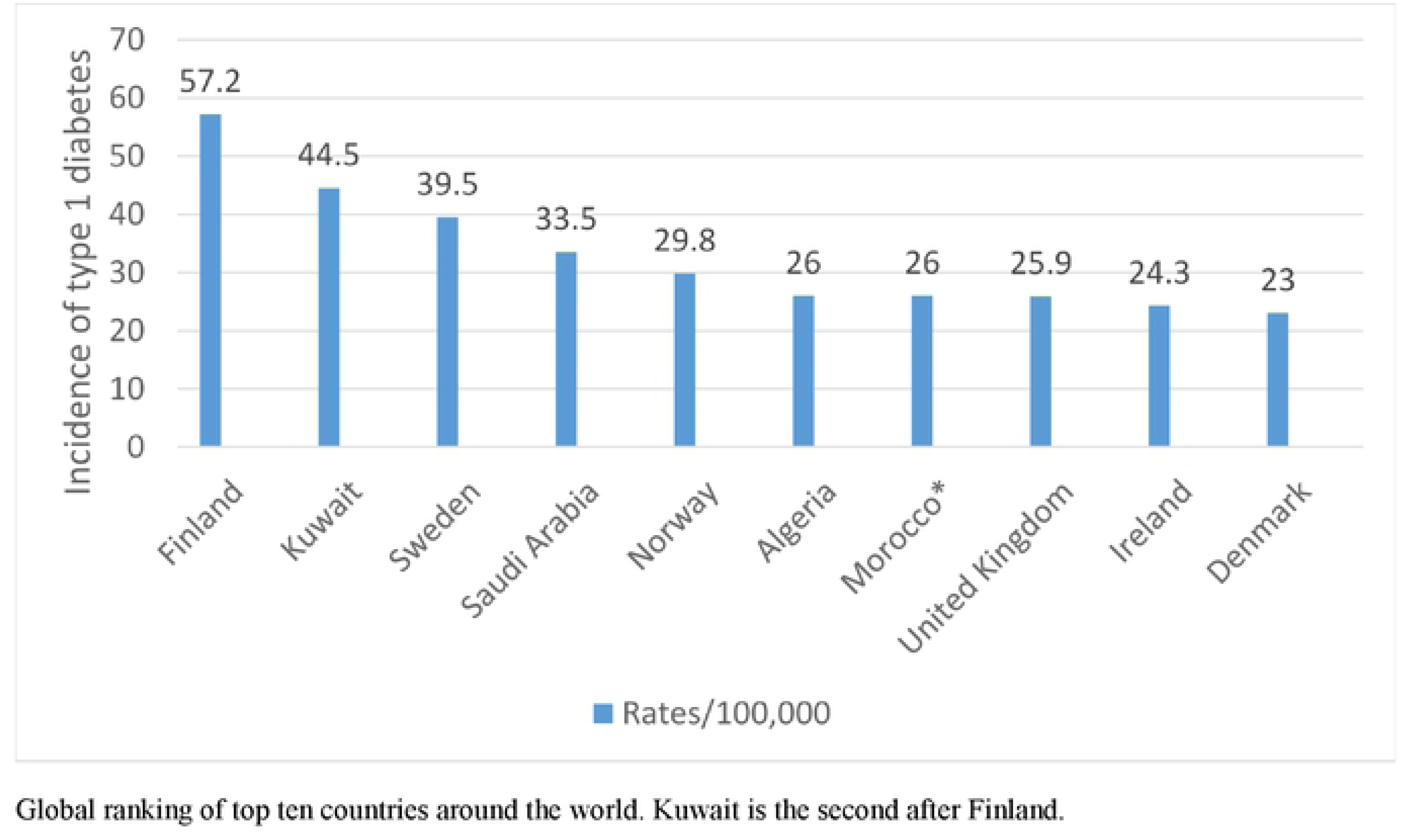
Top ten countries with highest rate of type 1 diabetes around the world.

T1D is defined as immune-mediated diabetes [3], which is caused by genes and environmental factors, such as viruses, that trigger the disease [4]. It is usually found in children, adolescents, and young adults, especially those with hyperglycemia and diabetic ketoacidosis [4]. The tendency to develop T1D, as shown in other autoimmune diseases, can be passed down through subsequent generations. Numerous studies, such as TrialNet, BABYDIAB in Germany, All Babies in Southeast Sweden [ABIS], Bart’s Oxford Family Study [BOX] in the U.K., and the Diabetes Autoimmunity Study in the Young [DAISY] in U.S.A., have focused on the causes of T1D and the possible ways to prevent or mitigate the disease.

Although a significant proportion of patients with T1D lack a family history of the disease, there is considerable familial clustering, with an average prevalence of 6% among siblings, compared to the 0.4% of the U.S. population. Moreover, there is a 3.8% risk of T1D among the Japanese siblings of patients with T1D, compared to the 0.01∼0.02% prevalence in the Japanese population [5, 6]. In this regard, the sibling ratio (λs) can be calculated as the ratio of the risk to siblings over the disease prevalence in the general population or λs = 6/0.4 = 15 and 3.8/0.01∼0.02 => 100 for the U.S. and Japan, respectively [5, 6].

Familial aggregation refers to the occurrence of a given trait shared by family members (or a community) that cannot be readily accounted for by chance. In this case, a family with a sibling or parent with T1D is much more likely to pass the disease on to other family members [5, 7]. The rising incidences of this disease in Kuwait might be due to rapid lifestyle changes, including “a sedentary lifestyle, changes in breastfeeding practices, and autoimmune deficiency caused by greater hygienic standards and low vitamin D levels, which are highly prevalent in the region in spite of the sunshine” [8]. Meanwhile, the rate of consanguinity and endogamous marriages in Kuwait is quite high at 22.5%–64.3% [9, 10]. Previous research has shown that localizing the root cause of complex diseases has been successful among such populations [11]. Kuwait Autoimmune Diabetes Study (KADS) is a familial case/control study. It aims to elucidate the islet autoantibodies profiling of Kuwaiti children and adolescents with T1D with T1D and their first-degree relatives. Recruiting families, including children, for clinical research studies can be challenging, and re-recruiting former participants can be even more difficult. Since Kuwait is known for its strong sense of familism, affiliative obedience, and filial obligation, a familial study of this disease may disclose certain causative agents responsible for passing the disease on to subsequent generations.

To the best of our knowledge, this is the first well-structured longitudinal familial study aimed at characterizing this serious disease. For this study, it provides general background information about the Kuwaiti national population, with its strong sense of familism, peculiar form of economic development, and history of pro-nationalist pressures associated with Arabic and Islam. Although the authors have extensive experience in familial recruitment in the U.K., the U.S., and Bahrain, there were certain difficulties in convincing, recruiting, and re-recruiting Kuwaiti families.

Several barriers to recruitment of participants were identified in literature. The attitudes of patients towards participation in research are considered as the main psychosocial barriers encountered in healthcare research [12, 13]. Distrust of outsiders, researchers, is one of the main psychosocial barriers encountered in recruitment for research [12, 14]. The inclusion of those who are insiders in the candidates’ community such as healthcare providers or community-based healthcare organizations may help in overcoming the mistrust barrier to study recruitment [12, 15]. Pervious researchers recruited via clinicians or nurses as they are more capable of gaining the patients’ trust [16], however, Sullivan-Bolyai et al deduced that physicians see recruitment for research as either job that consumes their time with no compensation or takes away time from patient care. Some researchers overcame this barrier by providing monetary incentives or offering other incentives such as purchasing a laptop, textbooks, journal or professional organization subscriptions, or sponsoring professional health care conference attendance [16]. Yet, these are costly and represent a burden on the budget of any study [17]. Families often recognize nurses as ‘trustworthy’, since the biggest trust barrier is lack of knowledge about the researchers, the healthcare research, and lack of trust in scientists [12]. Relationships represented an important human factor of recruitment; regular contact with patients and their families between visits may contribute to patients’ willingness to participate in research studies and positively affect the retention of participants [16]. Morgan et al highlighted that the presence of incurable illness reduced patients’ and families’ inclination to participate in research studies versus others with curable diseases who were more willing to participate [12, 18]. While Schutta et al deduced that the main drive for participation in research studies is the hope for a cure from the disease [19]. Therefore, the main goal is to contextualize the strategies and difficulties and to share our experiences with those interested in culturally oriented populations (e.g., Arabs).

## 2. Methodology

This study was approved by the Scientific Board of the Dasman Diabetes Institute (DDI). From November 2016 to September 2018, the authors conducted and modified several waves of research to recruit a sample of T1D patients, along with their first-degree relatives. In this case, the subjects were families with either an affected member or sibling or offspring (see Figure 2), with the goal of determining the T1D causing agent(s).

### 2.1 Physician-based approach

Since Kuwaiti families are generally obedient to their doctors, the authors decided to recruit the patients through their endocrine physicians. Relying on the physicians was helpful since they clearly explained the purpose of the study to the patients and their families. Health information management (HIM) personnel and the research coordinator (RC) were also part of the research team to facilitate the recruitment process.

### 2.2 Home visits

Making home visits was another approach for meeting the families’ requirements. In this case, a team consisting of one nurse, two phlebotomists (a male and a female, since some participants refused to be seen by the opposite gender), and a driver of the institute’s car was arranged. After receiving their consent, the list of families was categorized by the RC according to the districts/areas in which they lived. In order to prevent any duplication or multiple coding, the Dasman Diabetes Institute’s Biobank was responsible for label preparation and family coding. Two laboratory technicians were also available to process the samples as soon as they arrived.

### 2.3 Diabetes educator

Evidence-based medicine recommends educating patients about their respective diseases. In this regard, diabetes education has been shown to be effective in assisting patients to make informed decisions regarding the management of their disease [18]. In addition, such education can help reduce disease-related tension and improve an individual’s quality of life.

In this study, the patients were referred to a diabetes educator (DE), who taught them about the disease, how to improve their diabetes control, and how to manage their diabetes on a daily basis. The final and current mode of family enrollment was tailored according to the DE. The authors utilized this close relationship between the patients, parents, and DE to explain the importance of KADS, clarify the purpose of the study, and present the expected outcomes. Overall, the RC was in direct contact with the DE. Moreover, the RC was responsible for recording the data and the follow-up visits with the patients and their family members. Meanwhile, tubes and labels were necessary for categorizing the samples. Finally, the participants were requested to fast for approximately eight hours before their samples were collected.

## 3. Results

From November 2016 to September 2018, three main scenarios were performed. Each scenario was modified in order meet the needs of the family recruitment process. Overall, a reasonable number of families agreed to participate in the study. However, some individuals either did not respond to the RC or rejected or postponed their participation. Another drawback was that, according to Kuwaiti parental law, consent for minors under the age of 21 must be signed by the father and not the mother. Given the fact that the fathers rarely attend the routine visits to the physicians, it was hard to get the consent later via our RCs.

### 3.1 Physician-based approach

During physician-based approach, we collected 22 consented families to participate; however, none of them appeared. Despite our RC repeatedly contacting them, they still did not show any interest, Table 1. Moreover, when the physician retired, the team were faced by clear rejection of participation from the families.

**Table 1.**

| **Reasons for lack of success of physician-based recruitment approach** |
| --- |
| Linguistic slang |
| Lack of trust in healthcare research |
| Insufficient time of physicians to explain study purposes and requirements |

### 3.2 Home visits

Home visit approach was mainly proposed to tackle resistance to visit the institute for sample collection, and to avoid transportation of any disabled family members. It was likely that this flexible approach would increase participation rates, as logistical barriers such as inflexible work schedules or lack of transportation of disabled family member. Although a skilled and multidisciplinary team was arranged, this approach was not applied, due to the following reason (see Table 2).

**Table 2.**

| **The obstacles faced the home-visits approach** |
| --- |
| Both male and female phlebotomists were required |
| The availability of all family members |
| The fasting of more than 8 hours, given that the majority were children |
| The lack of private transportation for the follow-up visits by some of the team members |
| The time between sampling and laboratory processing |

### 3.3 Diabetes educator

At the DDI, all the patients were referred to the DE, who educated them and their respective families about diabetes management. In total, 31 (3.86% per month) Kuwaiti families with at least one affected member were successfully recruited (see Table 3).

**Table 3.**

| **The obstacles during the current phase** |
| --- |
| Arranging home visits to draw samples from the disabled members of the families |
| The inability of the participants who had school or work commitments during the weekdays |
| The difficulty of arranging all the family members in the same location |
| The inability to fast for more than 8 hours |
| The permission to leave school or work |
| The needle phobia of certain participants |
| The mistrust in diabetes-related research |
| Negative previous experience of healthcare research |

## 4. Discussion

In this article, we draw upon our own experience conducting a familial aggregation study. We explain the challenges we faced through our recruitment journey and how the research team overcame the challenges faced with initial recruitment strategies. Additionally, we report on our successful recruitment strategy which can be adopted by researchers in similar culture. From our experience, we faced some barriers which could be categorized into: psychosocial barriers and physical barriers.

### 4.1 Psychosocial barriers

The anti-research attitudes of patients towards participation in research are considered were the main psychological barriers. In our study, the DEs were challenged by tribal and religious beliefs of the candidate families. Participation in research was believed by some to be opposition of God’s fate and that all research was purposeless because they believed God is the only healer. Other families had negative attitude from some families during recruitment, as they believed no direct benefit to the affected case or their family, although the DEs often highlighted that the evidence and information gained from this research may help scientists and doctors to learn more about this condition and ultimately pave the road to find a cure or prevent that condition. On the other hand, some families had some unrealistic expectations of their participation, such as finding a cure for T1D. Our DEs frequently enforced the purposes and expected outcomes of the study, as it is crucial in gaining the participants’ trust and improving retention rate. Mistrust in the research system and negative experience from former participation in research was another psychosocial barrier. Some candidates claimed they were never informed with the results of the study in which they participated. Due to the strong familial structure of the Kuwaiti families, some candidates were afraid of leakage of data and being stigmatized as families with hereditary disease (T1D in our study) and others were in denial or fear of knowing that other controls from the family maybe susceptible or diagnosed with T1D in the future. Luckily, those candidates are provided with education by DEs within the same clinics where they were recruited. Continuous education was successful in most of the cases to change the candidates minds towards participation in the study. In addition, the candidates were assured about the confidentiality of the data and that no reference to their identities, families or tribes in any published article about this study. Mistrust of the healthcare research system and reliance on their traditional beliefs and religion and worship for healing were the main psychological barrier we experienced throughout our work. Relatively the same barriers have been raised by others as well [13]. These barriers were reduced by providing continuous education during our study period as found by previous studies [13].

The strong relationships and continuous support families received from DEs encouraged them to participate in our study, which was deduced by others as well [16].

### 4.2 Physical barriers

Physical barriers to recruitment in our study included: time constraints, health related barriers (such as having a disabled family members), distance, language barrier or linguistic slang and financial barriers. All of which were reflected on the intensity recruitment efforts, recruitment rate and retention rate.

Distance was not an issue for most of the participants in our study, as the study recruitment site was located at the diabetes healthcare institute, where the cases were treated/followed up. That increased the participation convenience for the subjects and reduced some of the physical barriers. However, transportation for disabled family member was an issue for some families. Thus, the study team proposed the home visit approach for sample collection to reduce the burden on the family. With enough staff and funds this could be a successful approach to tackle this barrier. However, this approach faced several logistic barriers in our study, and was not preceded, resulting in losing some candidates who had disabled family member. The team had to set-up a database to track the contact with the consented families and to convince them to participate and to schedule appointments for samples’ collection and to confirm these appointments, while working around the families’ schedule and the available staff’s schedule. Those challenges included; the shortage of staff of lab technicians and drivers, sensitivity of storing the samples in the weather conditions of Kuwait during the time from collecting to processing of the samples and working around the families’ schedule to arrange for the appointments. Language barrier was a main barrier, given the fact that all lab technician are non-Arabic speakers, as the subjects, especially children, would need assurance while taking the blood samples.

Time was the main physical barrier for recruiters and participants of our study. The treating physician’s limited time was a barrier against clear explanation of the study purposes and requirements. The alternative was utilizing the close relationship DEs had with families. DEs were Arabic speakers who follow-up closely with diabetic patients and their families through education clinic visits and telecare. Through these close relationships, DE had more time than physicians and can explain the purpose and requirements of the study in their language. Initially, the DEs would get the families’ consent and then forward the contact to the RC, who would finalize the logistics with the phlebotomy, the laboratory and the Biobank for samples’ collection, processing and storage, then would set up the appointments for sample collection. The problem which often faced the research team is that the families would postpone the appointment for sample collection due to work or school commitments, and ultimately declining participation. The research team brainstormed alternative ways to facilitate sample collection on the same day of signing the consent form. Finally, the team created a fast-track to finalize the logistics within the same time of the family’s visit to the DE clinic and take the consent and the samples on the same day to avoid any confliction with the family commitment and scheduling. Moreover, the future sample collection visits were rescheduled on same days with the follow-up visits with the DEs to relieve the burden on the family and the interruption of their work/school commitments. We accommodated most appointment time requests, excluding cases, who had to be fasting at least 8 hours prior to sample collection. This helped to address the scheduling barriers where some participants were difficult to obtain samples from due to time constraints.

## 5. Conclusion

T1D is a common, multifactorial disease with strong familial clustering. In Kuwait, the incidence of T1D among children aged 14 years or under is the second highest in the world, with the number of cases increasing approximately 2.4% per year. Annually, a significant part of the Kuwaiti healthcare budget is specified for the management of lifelong chronic diseases. Since T1D usually starts in early childhood, the affected member will be under severe stress, which negatively impacts his/her quality of life. Although most new T1D cases are sporadic, first-degree relatives have an increased risk of developing the same disease. Evidentially, the high rate of consanguinity and intermarital situations might have contributed to the epidemic growth rate of this disease. Therefore, examining KADS should be one of the highest priorities in diabetes-related research and healthcare.

Although the authors have thorough experience in recruiting T1D families, the present study is the result of 23 months of research regarding Kuwaiti T1D families with at least one affected member. It is important to note that, in Kuwait, convincing families with an affected member was an extremely difficult task, due to their strong sense of familism, linguistic slang, culture, and religious traditions. Throughout the above period, we observed that Kuwaiti families (or, possibly, families in general) tend to feel more relaxed if the person is accustomed to their culture and is comfortable using linguistic slangs. This might have been the reason behind our lack of success in our physician-based approach, as our physicians were Kuwaitis but were approached by a non-native RC. Similarly, the DE was used to the Kuwaiti culture and slangs because she was born in Kuwait. Of course, this attitude cannot be considered racist, rather it is the cultural approach and psychology of people. All the study participants had conflicting commitments, including work, schools, familial activities. Our study team had to work around the schedules of all family members, giving appointments for sample collection. To overcome that barrier, the study team managed to simplify the logistic barriers, including phlebotomy, laboratory and biobank logistics for sample collection, processing and storage. This way, the participants would be able to provide the samples on the same time they had consented during their visit to the DEs clinics. This flexible approach in means of participation relieved the RCs and encouraged participation of the families at relatively low costs. Scheduling families and confirmation of sample collection appointments using DEs telecare phone/WhatsApp, facilitated the continued study participation; and helped to overcome the barrier of timing.

### Lessons from the recruitment process

Throughout this 23-month journey, the authors came across certain obstacles when recruiting the culturally focused families. Committed research staff may be able to brainstorm alternative approaches to recruitment and logistic steps of sample collection. Thus, the current DE-oriented approach is a suitable scenario for recruiting families with strong familism beliefs. However, we could have prevented the loss of certain families if we had planned for family visits on, for example, open weekends and after hours. Moreover, it would have been more rewarding if we had provided the families with simple incentives such as nutritious meals. Linguistic slang is a critical factor in familial recruitment in Kuwaiti families.

### Limitations

The purpose of the present study was to characterize Kuwaiti Arab families. However, it disregarded Kuwaiti families with non-Arab mothers. This was simply due to genetic segregation, and it had nothing to do with racism. It narrowed our options in the recruitment process. Finally, it is important to note that there were several families that simply mistrusted diabetes research in general and refused to take part in this study or in any healthcare-related research.

## Acknowledgement

The authors would like to acknowledge the help of Ms Faten Sukkar for her initial support and conduction of the meetings. The authors would like to thank Mr. Sriraman Devarajan and his team for their cooperation and help in processing the samples and Mrs. Lorelee Tabada for her extended efforts in taking blood samples from KADS families’ members. The authors would like to thank Enago (www.enago.com) for the English language review.

## Authors’ contributions

MJ, as the principal investigator of KADS, designed and wrote the manuscript. NT revised the manuscript and contributed in the family recruitment process. ZR contributed extensively in the family recruitment process, while HA collected the data and coordinated with the families. MA, MM, FA, and HA served as the treating physicians.

## Declaration

The authors have no affiliations with or involvement in any organization or entity with any financial interest

## Ethical Approval

The current research was conducted after obtaining written approval from Dasman Diabetes Institute Ethical Review committee, RA 2016-015, and informed consent was attained by each participant in written.

